# An exactly solvable, spatial model of mutation accumulation in cancer

**DOI:** 10.1101/084038

**Authors:** Chay Paterson, Martin A. Nowak, Bartlomiej Waclaw

**Affiliations:** SUPA, School of Physics and Astronomy, The University of Edinburgh, Mayfield Road, Edinburgh EH9 3JZ, United Kingdom; Program for Evolutionary Dynamics, Harvard University, One Brattle Square, Cambridge, Massachusetts, USA.; Department of Mathematics, Harvard University, One Oxford Street, Cambridge, Massachusetts, USA.; Department of Organismic and Evolutionary Biology, Harvard University, 26 Oxford Street, Cambridge, Massachusetts, USA.; Centre for Synthetic and Systems Biology, The University of Edinburgh

## Abstract

One of the hallmarks of cancer is the accumulation of driver mutations which increase the net reproductive rate of cancer cells and allow them to spread. This process has been studied in mathematical models of well mixed populations, and in computer simulations of three-dimensional spatial models. But the computational complexity of these more realistic, spatial models makes it difficult to simulate realistically large and clinically detectable solid tumours. Here we describe an exactly solvable mathematical model of a tumour featuring replication, mutation and local migration of cancer cells. The model predicts a quasi-exponential growth of large tumours even if different fragments of the tumour grow sub-exponentially due to nutrient and space limitations. The model reproduces clinically observed tumour growth times using biologically plausible rates for cell birth, death, and migration rates. We also show that the expected number of accumulated driver mutations increases exponentially in time if the average fitness gain per driver is constant, and that it reaches a plateau if the gains decrease over time. We discuss the realism of the underlying assumptions and possible extensions of the model.

## Introduction

Every cancerous tumour is initiated by a single rogue cell which has accumulated genetic or epigenetic alterations which enable it to proliferate faster than cells from normal tissue. This process is stochastic and occurs with a surprisingly small probability given the frequency of cell division in the human body [1]. However, once initiated, the progeny of the first neoplastic cell will likely eventually acquire further alterations. Some of these alterations are benign and constitute what are called “passengers”; others, called “driver mutations”, may further increase proliferation, usually by decreasing the responsiveness of cells to signals which normally halt division. Cancer is thus governed by the rules of biological evolution: driver mutations endow cells with selective advantage, whereas passenger mutations behave as neutral mutations “hitchhiking” on clonal expansions fuelled by the drivers. It is believed that 3-15 drivers are present in a majority of cells in a clinically detectable tumour [2–6].

The possibility the accumulation of new drivers during cancer progression raises a number of interesting questions: How is the growth of a three-dimensional mass of cells affected by drivers? How are drivers distributed in the tumour? How many drivers are expected in a tumour of a given size or age? Answering these questions is important in order to understand the observed cancer incidence rates [1, 6, 7], predict how many oncogenes and tumour suppressor genes are yet to be found [8], and to explain the observed levels of genetic heterogeneity in tumours [9–12].

Both initiation and progression have been modelled mathematically [13–17], and many theoretical and computational models have already addressed the above questions. “Well-mixed” models [8, 18–21] which omit all spatial structure show that new mutations accumulate faster over time [8, 21]. This process has been also studied in spatial models [22] in which the tumour is a single ball of cells whose radius grows linearly in time, and in more idealized, lattice models [23–27].

Recently, computational models have been proposed [12, 28, 29] in which cancerous tumours are conglomerates of one large primary tumour with many surrounding microlesions, presumed to form when cancer cells detach from the primary lesion and resume growth at an adjacent site. This is similar to metastasis – the ability to invade and form new microlesions elsewhere in the body – which is arguably the defining characteristic of cancer: however, these models [12, 28, 29] are also applicable to short-range invasiveness, perhaps as short as a few micrometers. In fact, the presence of cancer cells in regions surrounding the primary lesion have long been used as a predictor of cancer recurrence [30, 31] following surgical excision of solid tumours.

The movement of cells in such models is understood as caused by cells undergoing epithelial-to-mesenchymal transition and local invasion[32], or possibly “sprouting” due to mechanical instability [33, 34], though the models tend to be neutral as to the underlying mechanism. Having left the primary lesion, cells enjoy a better access to nutrients and oxygen and can proliferate faster. The tumour has a complex structure which enables faster expansion than if it were made of a homogenous and simple cluster of cells. This is distinct from models of metastasis [35, 36] which assume spatially separated, distant lesions.

In this work we study a simplified, mathematical counterpart of the models [12, 28, 29] that can be solved exactly. This enables us to study tumours of any size, including clinically relevant sizes. Our model belongs to the class of age-structured population models, which were first used in population ecology and epidemiology [37, 38], but have more recently been applied to modelling metastasis in cancer [35, 39].

Here, we substantially extend these earlier models by including many cell types, each corresponding to a genotype with a new driver and hence a different growth rate. Surprisingly, our extended model can still be analytically solved in several important cases, allowing us to formulate hypotheses as to how rapidly driver mutations accumulate in cancer.

## The model

We model the tumour as a collection of microlesions made of cancer cells (Fig. 1A). Microlesions do not interact with one another; this is crucial for the analytic solubility of the model. This rather strong assumption could be justified by interpreting microlesions as spatially-separated metastases: however, even if microlesions are fragments of the same primary tumour, numerical simulations (Ref. [12], Extended Data) indicate that as long as microlesions are separated by normal tissue, their growth rate is only minimally affected.

**Figure 1:**
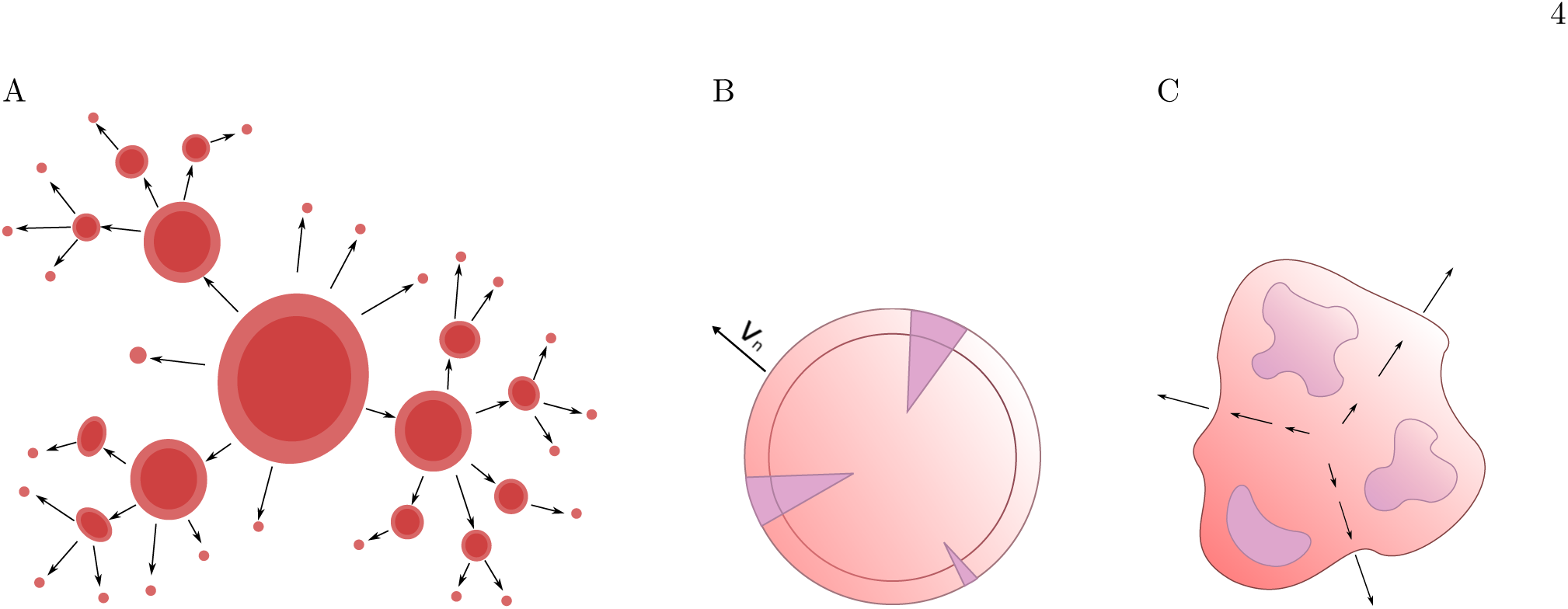
A: The model assumes that the tumour is made of discrete microlesions. Cells migrate from microlesions (arrows) and establish new microscopic lesions. All lesions increase in size over time. B,C: two different growth models of individual microlesions. In the surface growth model (B), cells replicate only in a narrow layer of constant thickness near the surface, and the radius of the lesion increases with velocity *v*_*n*_. In the volumetric growth model (C), replication occurs everywhere and the microlesion grows exponentially which causes the whole lesion to “inflate”. Arrows show the expansion velocity which is small close to the centre and increases towards the surface. Purple areas correspond to new mutations.

Each microlesion is fully characterized by two numbers: its age *a* and its type *n* which describes the genetic makeup of the lesion. For simplicity, we take *n* to be the number of driver mutations in the cell which initiated the lesion. This convention does not need all microlesions to be genetically the same, but only for their evolution to be completely determined by the two variables {*n*, *a*}. In what follows we shall assume that lesion {*n*, *a*} contains at most two different genotypes, *n* and *n* + 1, with relative abundances *r*_*n*_(*a*) and 1 − *r*_*n*_(*a*), respectively. Note that we do not model individual cells explicitly: rather, we model only the behaviour of microscopic lesions which contain thousands or even millions of cells.

The dynamics of the model is governed by three processes: growth, the emergence of new drivers caused by mutations, and the creation of new microlesions by migration.

*Growth.* A microlesion {*n*, *a*} grows deterministically and its volume at age *a* is given by *V*_*n*_(*a*). The proportion of the *n* + 1-th genotype also grows deterministically and equals 1 − *r*_*n*_(*a*). The functions *V*_*n*_(*a*) and *r*_*n*_(*a*) depend on the growth model of a single, isolated lesion and we shall specify them later (Results, “Multiple driver mutations”, equations (30) onwards). The spatial character of the model manifests itself in our choice of these functions; for example *V*_*n*_(*a*) is very different for three-dimensional growth in which replication occurs only on the surface, and “well-mixed” growth in which growth occurs in the whole volume and all cells replicate with identical rates. This is the only component of the model affected by spatial arrangement of cells.

*Driver accumulation.* Cells in microlesion {*n*, *a*} can gain a new driver *n*+1 with some small but non-zero probability. This represents the occurrence of genetic mutations and is accounted for in the form of *r*_*n*_(*a*), the fraction of cells that do not contain the new *n* + 1-th driver. Therefore, a lesion changes its genetic composition in a deterministic way in this model.

*Migration.* A lesion {*n*, *a*} seeds new microlesions with rate *ϕ*_*n*_(*a*). A new lesion has age *a* = 0 and, with probability *r*_*n*_(*a*), it inherits the same type *n* as the parent lesion, or with probability 1 − *r*_*n*_(*a*) it is assigned type *n* + 1. This accounts for what would have happened if we modelled individual cells within the lesion, and cells of different types migrated from the lesion with probabilities proportional to their abundances. For example, the probability that a cell of type *n* +1 migrated and established a new lesion of type *n* +1 would be equal to the fraction 1 − *r*_*n*_(*a*) of cells of type *n* + 1 in the lesion.

The model defined by the above postulates is stochastic: while growth and mutation are deterministic, migration (creation of new lesions) is a random process. In what follows we shall focus on the analysis of the mean behaviour of the model, averaged over many realisations of the stochastic process, and thus we shall ignore fluctuations. The role of fluctuations will be discussed in Section *Comparison to simulations.* Let *f*_*n*_ (*a*, *t*) be the expected number of microlesions of age *a* at time *t*. The time evolution of *f*_*n*_(*a,t*) is governed by the following two equations,

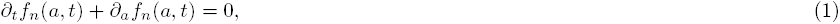

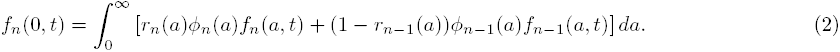

with the initial condition *f*_*n*_(*a*, 0) = *δ*_*n*,1_*δ*(*a*) in which *δ*_*n*,1_ is the Kronecker delta and *δ*(*a*) the Dirac delta function. Equation (1) describes the shift of the distribution in time - the “ageing” of the microlesions. Equation (2) describes the birth of new lesions (through migration) and can be thought of as a boundary condition for Eq. (1). The function *V*_*n*_(*a*) which defines the growth of a single lesion does not appear in these equations, but it appears in the equation for the total volume of the tumour,

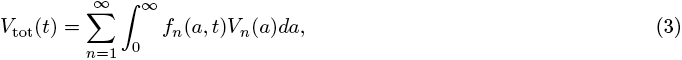

and the average number of drivers per cell,

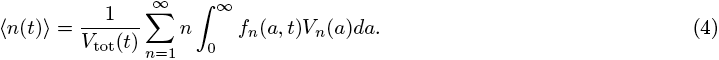

Although equations (1, 2) are quite general, their hierarchical structure allows an exact formal solution (Methods, “General solution with multiple types of microlesions”, equation (53) onwards). To model cancer in this framework, we must define the functions *r*_*n*_(*a*),*V*_*n*_(*a*), *ϕ*_*n*_(*a*) such that they describe the growth of individual microlesions.

## Results

### Only one driver

We first consider three special cases of the model with only one type of microlesions. This will form a set of “null models” against which we will compare the multiple-driver models from the next section.

#### General solution of Eqs. (1,2)

In the case of one type of cells (no new driver mutations), *r*_1_(*a*) = 1 and *r*_*n*_(*a*) =0 for *n* > 1, and Eqs. (1, 2) reduce to the McKendrick-von Foerster equation without removal [38]:

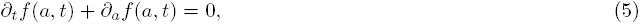

with the boundary condition

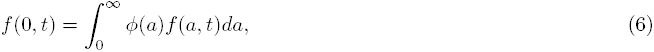

and the initial condition *f*(*a*, 0) = *δ*(*a*). Note we have dropped the index “ 1” in *f*_1_(*a*, *t*) and *ϕ*_1_(*a*) for brevity. Equation (5) has a solution of the form *f*(*a,t*) = *F*(*t* − *a*)Θ(*t* − *a*) + *δ*(*t* − *a*), where *F*(*z*) is some non-negative continuous function, Θ(z) is the Heaviside step function, and the Dirac-delta term *δ* (*t* − *a*) accounts for the initial condition. The boundary condition (6) implies that

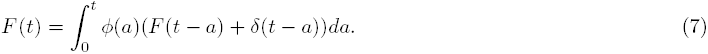

Laplace-transforming the above equation gives

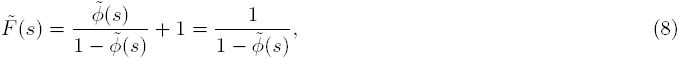

where the constant term 1 is due to the singular initial condition, and the functions marked with tildes are Laplace transforms: 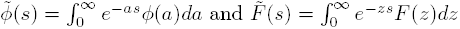.

Equation (8) enables us to calculate the distribution *f*(*a,t*), provided we can invert the Laplace transform. We shall see that this is possible in some important special cases (Results, “Multiple driver mutations”). However, even if the full distribution cannot be obtained, Eq. (8) provides us means to infer large-*t* behaviour of *f*(*a,t*). We note that 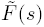 from Eq. (8) becomes singular when 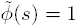 and behaves as 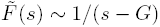 near any of the roots *G* of the equation

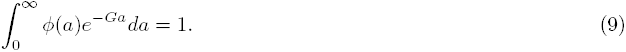

The inverse transform *F*(*z*) of 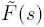 will thus contain terms ~ exp(*Gz*). One of these terms corresponding to the largest *G* dominates the long-time behaviour, and the distribution in this limit reads

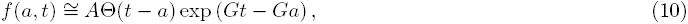

where *A* is some positive constant. Equation (9) is called the Euler-Lotka equation [37, 38] and its largest root *G* gives the (exponential) growth rate of the number of microlesions *N*(*t*) in our model as follows from

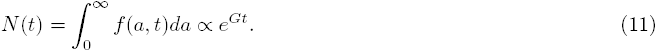

Similarly, the total volume (3) also increases exponentially,

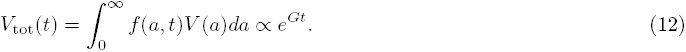

These results are well known in the literature of mathematical ecology. Here they were re-derived using a different approach which can be adapted to more complicated cases with many cell types (new driver mutations). We shall now use these general results to investigate three special cases of cancer growth.

#### Surface growth

We assume that microlesions are spherically symmetric, cells replicate only on the surface, and that the centre of the lesion is static. Let the volume (number of cells) of the lesion increase with its age *a* as

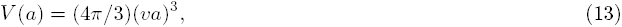

where *v* is the speed (in cells/day) with which the radius of the ball of cells expands in time. We further assume that cells can migrate only from the surface, thus

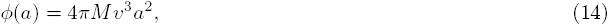

where *M* is the dimensionless migration probability representing the fraction of cells on the surface which escape and go on to eventually seed new microlesions. Taking the largest root of Eq. (9) we find the asymptotic growth rate *G* to be

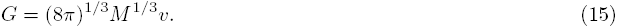

In fact, for this simple model we can find the exact expression for the number of microlesions *N*(*t*) and the total volume *V*_tot_(*t*). The Laplace transform of *ϕ*(*a*) reads 8*πMv*^3^/*s*^3^ and from Eq. (8) we obtain

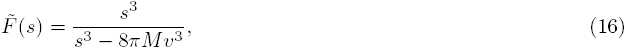

which can be inverted:

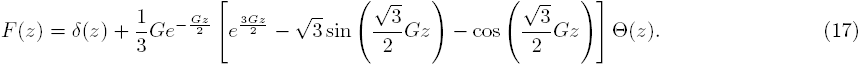

Here *G* is the growth rate from Eq. (15) and Θ(*z*) the Heaviside step. We thus obtain that the total number of microlesions increases in time as

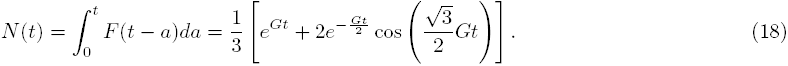

As expected, *N*(0) = 1 and *N*(*t*) ≅ (1/3)*e*^−*Gt*^ for *t* → ∞. The total volume (number of cells) is

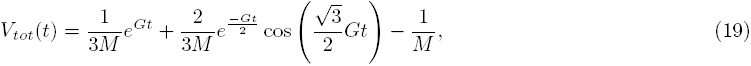

which also increases exponentially with time. Figure 2A shows how the total volume increases in time for a fixed *M* and a range of *v*, whereas Fig. 2B shows the same quantity for fixed *v* and different *M*. It is evident that in order for the tumour to grow to 10^11^ − 10^12^ cells (10-100cm^3^) over the period of 15 years which is thought to be typical for solid cancer, both *M* and *v* must be sufficiently large, otherwise there is not enough migration from the surface and the tumour grows sub-exponentially. On the other hand, *M*, *v* do not have to be very big: if *M* = 10^−6^ (only one cell per million migrates) and *v* = 0.1 (microlesion radius grows by 0.1 cell per day), a tumour can grow to a macroscopic size in a realistic time frame.

**Figure 2:**
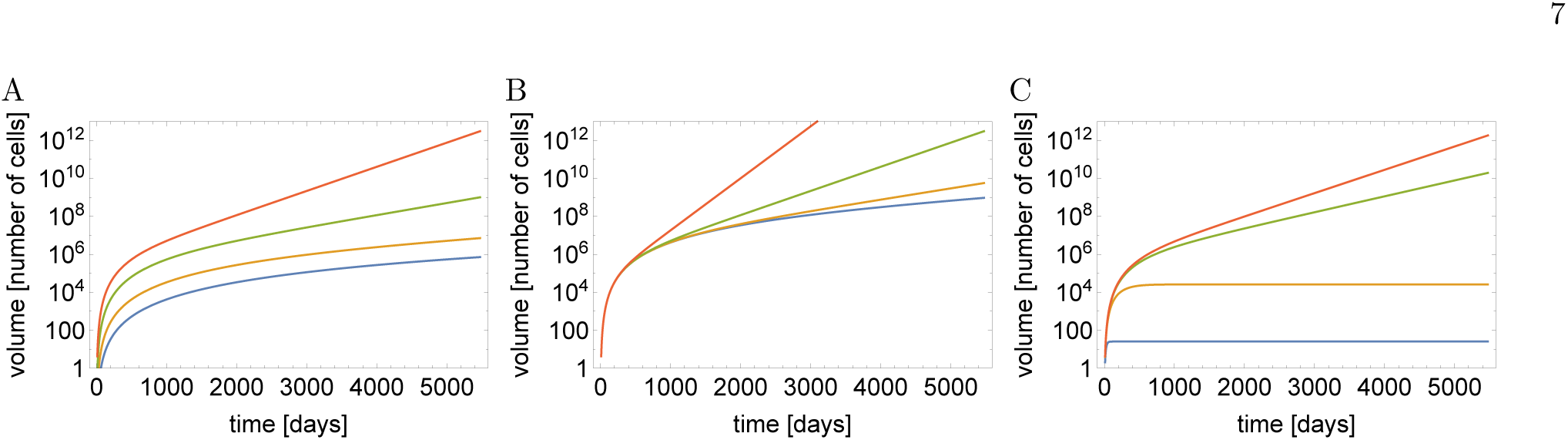
A: total tumour volume *V*_tot_(*t*) versus time for different single-lesion expansion velocities *v* = 0.01 (blue), *v* = 0.02 (yellow), *v* = 0.05 (green) and *v* = 0.1 (red), and the same migration probability *M* = 10^−6^. B: *V*_tot_(*t*) for *v* = 0.1 and four different *M* = 10^−8^,10^−7^,10^−6^,10^−5^ (blue, yellow, green and red, respectively). C: *V*_tot_(*t*) for the slow-down model, *M* = 10^−6^, *v* = 0.1 and four different *λ* = 0.1, 0.01,10^−3^,10^−4^ (blue, yellow, green and red).

#### Surface growth with decreasing replication rate

In reality, a lesion cannot be expected to grow continuously. Growth eventually slows down due to spatial constrains and limited nutrient supply which is why many tumours never reach detectable sizes [40, 41]. Let us consider a simple case of an exponential slow-down:

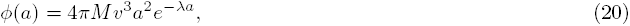

where *λ* > 0 is the characteristic time scale of growth decrease. As before, we assume that proliferation occurs only on the surface, and therefore

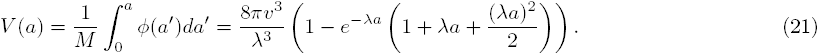

This function has a sigmoid shape and saturates at *V*(∞) = 8*πv*^3^/*λ*^3^. The formulas for *ϕ*(*a*), *V*(*a*) reduce to Eqs. (13,14) when *λ* = 0 i.e. when there is no slow-down. We can now calculate 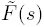:

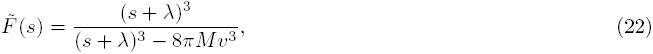

and, upon inverting the Laplace transform, we have

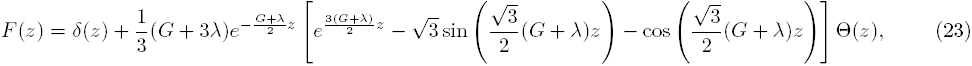

where

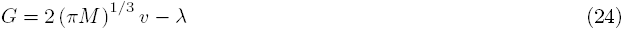

is the exponential growth rate of the tumour. We note that *G* > 0 only for sufficiently large *M*, thus the tumour made of microlesions whose growth slows down can grow exponentially as a whole only if sufficient migration is present. We can also calculate *N*(*t*) and the total volume exactly but since both expressions are rather lengthy and not particularly illuminating (SI Mathematica notebook), we quote here only the leading terms:

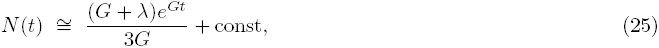

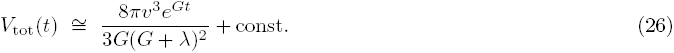

Figure 2C shows that the volume increases exponentially only if *λ* is small enough, otherwise growth becomes arrested after some time.

#### Volumetric growth

Finally, we shall consider the case in which all cells from the lesion are able to replicate and migrate. If cells replicate with rate *b* and migrate with rate *M*, we obtain

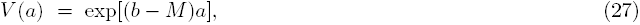

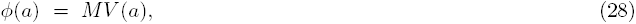

thus individual lesions grow exponentially in time. As expected, Eq. (9) predicts that the whole tumour grows exponentially with rate *G* = *b* i.e. the same rate as the birth rate of individual cells.

### Multiple driver mutations

We shall now extend these results to the case in which new drivers can emerge during tumour growth. Each new driver mutation speeds up growth as a result of increased proliferation/decreased death of cells.

#### Surface growth

We assume that each new driver mutation increases the expansion speed of microlesions. This can be motivated as follows: replication and death balance exactly for normal cells to ensure homoeostasis. A driver mutation increases the birth rate or decreases the death rate and hence the net growth rate becomes positive [42]. In the case of surface growth of a solid lesion considered here this net growth rate is assumed to translate into a radial expansion of the lesion with speed *v*_*n*_.

We first take the case *v*_*n*_ = *nv*_1_, i.e., each new driver contributes equally to *v*_*n*_. We have

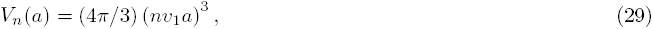

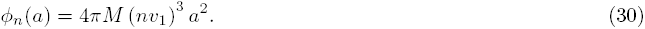

So far we have not specified *r*_*n*_(*a*), the fraction of cells with *n* drivers in the lesion of type *n*. To simplify calculations, we shall take

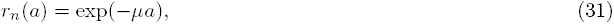

with some positive constant *µ.* The above equation can be rationalized if we assume that cells on the surface of the lesion mutate from *n* to *n* +1 drivers with a very small rate *µ* and that their selective advantage can be neglected. The dynamics of type *n* +1 in a lesion of type *n* is thus effectively neutral. A mutant sector that arose at time *a*_0_ will then have approx. (*a/a*_0_)^2^ cells on the surface. The fraction 1 − *r*_*n*_(*a*) of mutant cells obeys then the following equation:

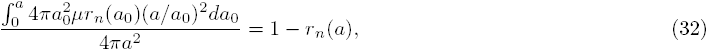

which can be solved yielding Eq. (31). However, for the assumed *v*_*n*_ = *nv*_1_ the selective advantage of type *n* +1 over type *n* is (*n* + 1)/*n* = 1/*n* > 0 and therefore our assumption about the population dynamics being neutral does not hold for small *n*. The dynamics of non-neutral mutant sectors is quite complex [22, 43] and leads to formulas for *ϕ*_*n*_(*a*)*r*_*n*_(*a*) which cannot be Laplace-transformed analytically. We therefore stick to a much simpler but perhaps less realistic Eq. (31). In Section “More realistic surface fractions *r*_*n*_(*a*)*”* we shall discuss *r*_*n*_(*a*) that is more biologically plausible while it still leads to analytically solvable models.

In Methods (“Surface growth with many drivers”), it is shown that the exponential growth rate of the number of microlesions of type *n* is

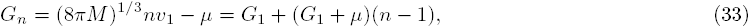

where *G*_1_ = (8*πM*)^1/3^*v*_1_ − *µ*. The growth rate can thus become negative for sufficiently large mutation rates *µ*. This effect is similar to what was found in the preceding subsection (“Surface growth with decreasing replication rate”). However, in contrast to this earlier result in which replication slowed down over time, resulting in fewer cells able to migrate and establish new microlesions, here the cells only change their type while their replication and migration is not affected. The total number of cells thus increases over time; we shall show this explicitly later in this section. A negative *G*_*n*_ only means that a subpopulation of type *n* cannot grow exponentially at long times: once a strain with *G*_*n*_ > 0 emerges, it rapidly dominates the population.

The volumes of cells of type *n*, *V*_tot,*n*_(*t*), as well as the total volume *V*_tot,*n*_(*t*) can be obtained analytically (Methods, “Surface growth with many drivers” and Supplemental Material). Figure 3A shows example curves *V*_tot,*n*_(*t*). The asymptotic behaviour of each subpopulation *V*_tot,*n*_ ~ exp(*G*_*n*_*t*) is clearly visible. Since *G*_*n*_ increases with *n*, microlesions containing more driver mutations grow faster and their population eventually overtakes that of microlesions with fewer drivers. For this reason, the total volume *V*_tot_(*t*) obtained by summing up all *V*_tot,*n*_ (Eq. (3)) increases faster than exponentially, as is plainly visible in the plots (Fig. 3B). In the large-*t* limit, we can show (Methods) that the volume behaves asymptotically as

**Figure 3:**
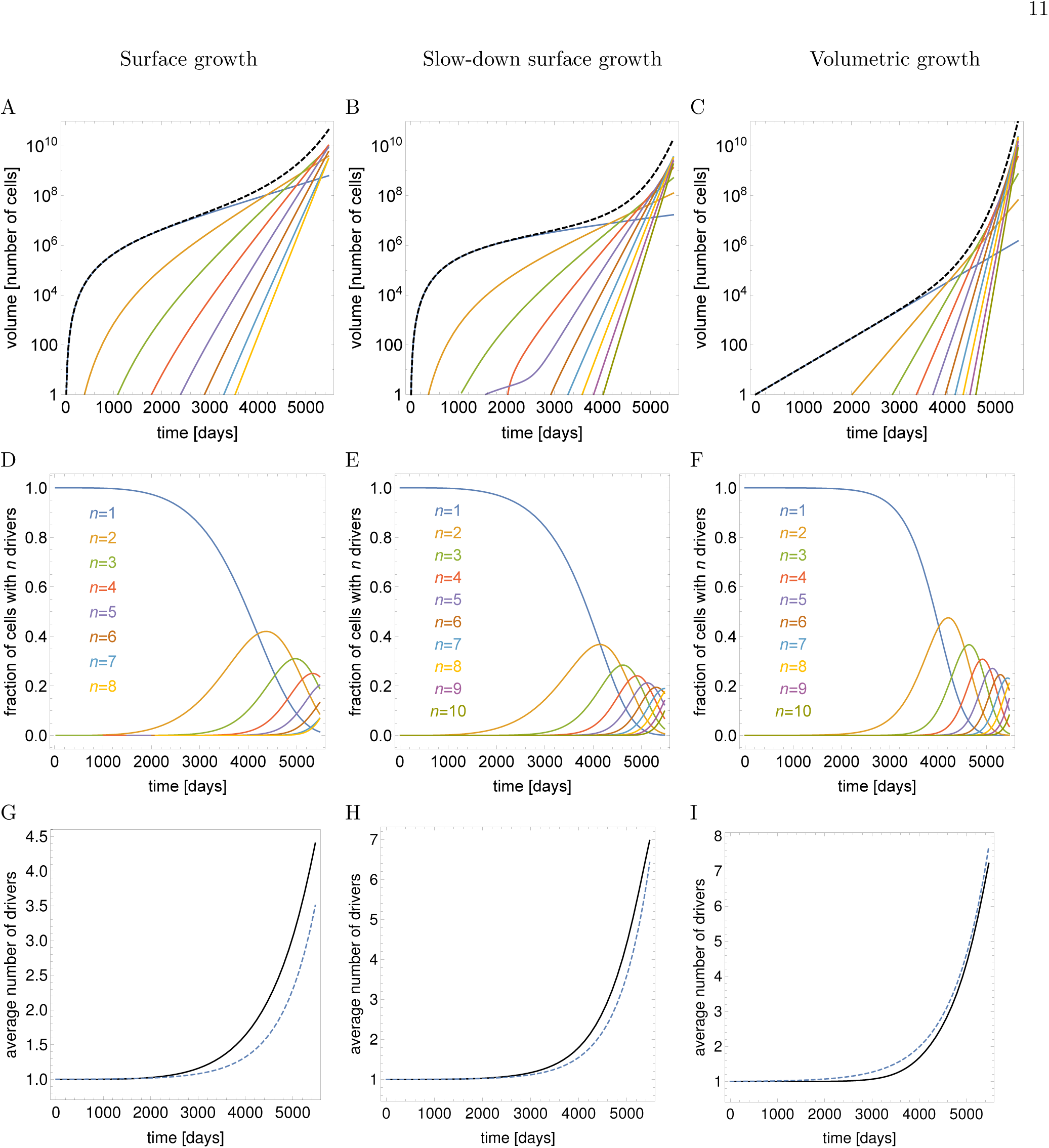
Plots of different quantities (rows of panels) characterizing the tumour as a function of time (days), for different growth scenarios (columns of panels). A,B,C: the total volume (number of cells) of microlesions with *n* driver mutations (coloured curves) and the total volume *V*_tot_ of the whole tumour (black dashed curve). D,E,F: the fraction of cells with *n* = 1, 2, 3,… driver mutations in the whole tumour (colours as in A,B,C). G,H,I: the average number of drivers 〈*n*(*t*)〉: exact calculation (black) and asymptotic formula (dashed blue). The columns are as follows. Panels A,D,G: from the section on the “Surface growth” model with parameters *v*_1_ = 0.0475, *M* = 10^−6^, *µ* = 2 · 10^−5^ (all rates in units day^−1^). The parameters *v*_1_,*M* have been chosen such that the tumour reaches about 10^11^ cells in 15 years (~ 5500 days) and accumulates 3 – 4 drivers, *µ* was assumed to be the same as in Ref. [8]. Black dashed curve in A is the total volume *V*_tot_ derived from Eqs. (3) and (57) with *F*_*n*_(*z*) given by (61). Solid black curve in G is the exact analytical calculation from Eqs. (4) and (61), blue curve the asymptotic approximation (35). Panels B,E,H: corresponding to the section on the “Surface growth with decreasing replication rate” model) for *v*_1_ = 0.053, *M* = 10^−6^, *µ* = 2 × 10^−5^, *λ* = 10^−3^. Blue curve in panel H is the asymptotic approximation (37). Panels C,F,I: volumetric growth for *b* = 0.0026, *M* = 10^−5^, *µ* = 2 · 10^−5^. Solid black curve is the exact analytical calculation derived from Eqs. (3) and (57) with *F*_*n*_(*z*) given by (77). Blue curve is the asymptotic approximation derived from (41). which turns out to be identical to Eq. (35) when we observe that *G*_1_ from Eqs. (36) and (33) differ only by *λ*. This means that the average number of drivers increases approximately exponentially over time with a rate that is not affected by growth of individual microlesions slowing down with age, and is the same for all λ. Example growth curves *V*_tot,*n*_(*t*) and the average number of drivers are presented in Figure 3B,E,H.

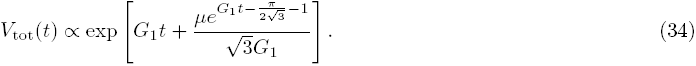

We can also calculate the average number of drivers (SI Mathematica notebook). Figure 3C shows that ⟨*n*(*t*)⟩ is small until late times when it rapidly increases. In fact, in the limit *t* → ∞ TO it can be shown (Methods) that

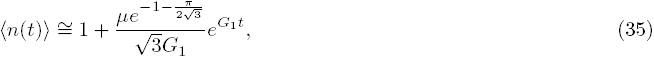

so the number of drivers accumulates exponentially over time. Note that the asymptotic formulas derived here are quite accurate even for finite times characteristic for tumour growth.

#### Slow-down surface growth

If growth of individual lesions slows down over time (as in “Surface growth with decreasing replication rate”), it can be easily obtained that

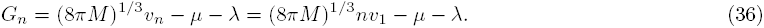

Note that the differences *G*_*n*+1_ − *G*_*n*_ do not depend on *λ* and are the same as in the case without slow down (Results, “Surface growth”). We shall see that this causes driver mutations to accumulate at the same rate as before. The average number of drivers reads (Methods)

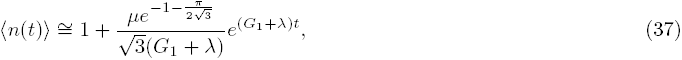

#### Volumetric growth

In the case of volumetric growth i.e. cells replicating in the whole volume of the lesion, we have

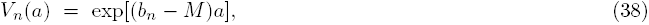

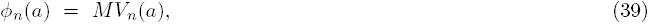

where *b*_*n*_ = *nb* is the net replication rate of cells with n drivers. We assume that *r*_*n*_(*a*) = exp(−*µa*) exactly as for the surface growth models (31). This formula correctly describes the proportion on cells of type *n* only if type *n* +1 does not have a selective advantage. In the case of selective advantage we are dealing with here, the formula is only approximately correct, as detailed in Section “More realistic surface fractions *r*_*n*_(*a*)”.

In the limit of long time we obtain (Methods) that the total volume and the average number of drivers behave asymptotically as

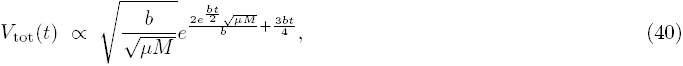

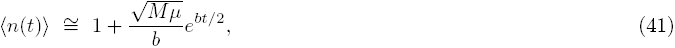

so that the total volume increases faster than exponentially, and drivers accumulate exponentially, similar to the surface growth models from previous sections. We can also compute exact expressions for *V*_tot_(*t*) and 〈*n*(*t*)〉 (Methods). Figure 3C,F,I shows the volume and the number of driver mutations as a function of time. Note that to reach *both* the same size as in the surface growth model (of 10^11^ cells) *and* a reasonable number of drivers (more than two), the net increment *b* of the growth rate per driver has to be very small.

### Driver mutations with decreasing selective advantage

In all previous cases, virtually all drivers accumulated at a late stage during the growth of the tumour. In this section we shall study the case when the selective advantage of the first three drivers is much bigger than that of all subsequent drivers. Statistical analysis of cancer incidence rates suggests that this is the case for some lung and colorectal cancers [44]. We take the surface growth model with the following *v*_*n*_:

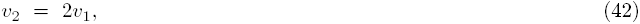

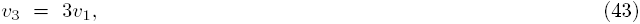

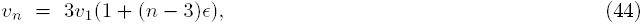

with some *v*_1_ and a small *ϵ* ≪ 1. The expansion speeds increases fast for *n* = 2, 3 and slow for *n* ≥ 4 drivers. Figure 4 shows that now most drivers accumulate earlier, before the tumour reaches a detectable size (*V*_tot_ < 10^9^). This causes the final tumour to become much less heterogeneous because almost all cells have the same driver mutations. Moreover, the total volume grows approximately exponentially for long times, rather than super-exponentially as in the previous models. These results are in agreement with recent experimental evidence on intra-tumour heterogeneity in colorectal cancer [45].

**Figure 4:**
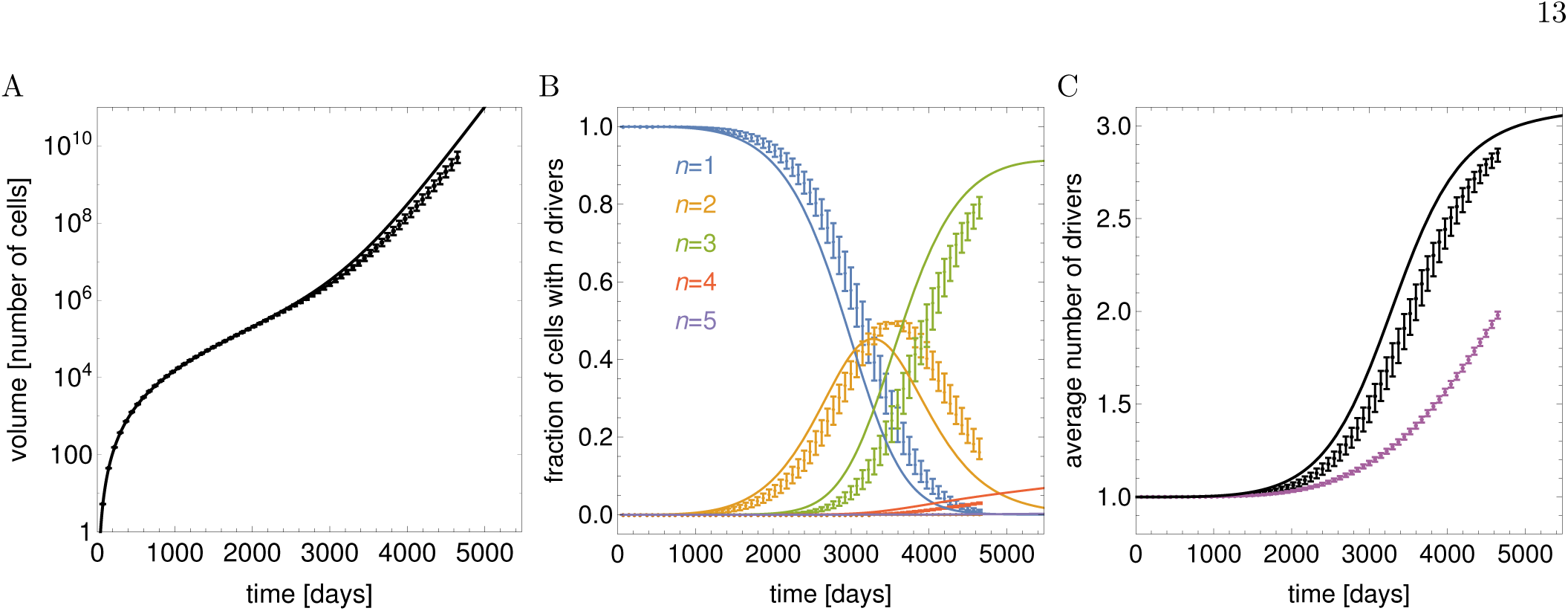
The model with three strong drivers as in Eqs. (42-44). A: The total volume of the whole tumour (black curve). B: the fraction of drivers with *n* mutations. C: the average number of drivers (black curve). In all cases *v*_1_ = 0.015, *M* = 10^−4^, *µ* = 2·10^−5^, *ϵ* = 0.01. Data points with error bars in all panels come from stochastic simulations of the original model (Section *Computer simulations*, for which 500 replicates were performed). In panel C, two sets of data points are presented. Purple points have been obtained by calculating the average number of drivers for each simulated tumour and then taking the average over many tumours. Black points represent averaging the fractions of cells with n drivers over many tumours (as in panel B), and using this to calculate the average *n*.

### More realistic surface fractions *r*_*n*_(*a*)

So far we have assumed the fraction of cells with *n* drivers to be *r*_*n*_(*a*) = exp(−*µa*) (see Eq. (31)). This simplifies calculations, but as already mentioned is not very realistic. In particular, as it is independent of *n*, Eq. (31) cannot take into account the expansion of the subpopulation of cells with *n* +1 drivers due to their faster growth. We shall now derive a more realistic form of *r*_*n*_(*a*) which is still simple enough to lead to analytic results for the total volume and the average number of drivers.

Recall that *r*_*n*_(*a*) is interpreted as the fraction of cells of genotype *n* in a type-*n* lesion that are able to migrate and establish new lesions. In the volumetric growth model this would correspond to all cells in the lesion that do not have the *n* +1 driver, while in the surface growth model only cells that are on the surface contribute to *r*_*n*_(*a*).

Let us first consider the volumetric model. If *b*_*n*_ is the growth rate of type-*n*, the fraction 1 − *r*_*n*_(*a*) of new mutant cells obeys the following equation:

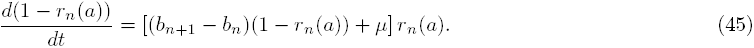

The first term (*b*_*n*+1_ − *b*_*n*_)(1 − *r*_*n*_(*a*))*r*_*n*_(*a*) corresponds to the rate with which the population with *n* +1 drivers increase due to selective advantage, and the second term *µr*_*n*_(*a*) accounts for new mutations. Equation (45) is of logistic type and can be solved yielding

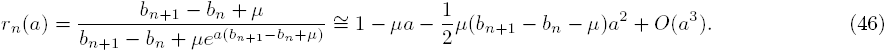

We can see that for small *a* the above expansion agrees with that of Eq. (31) which is the approximation we used before: the two formulae for *r*_*n*_(*a*) can therefore be expected to agree for *a* < *µ*^−1^. Although the exact form of *r*_*n*_(*a*) from the above formula leads to a Laplace transform of *F*_*n*_(*t*) (Methods) that cannot be inverted analytically, the Taylor expansion can be used instead; in the Supplementary Mathematica notebook we show example formulas for *V*_tot_(*t*) for this case.

We shall now consider the surface growth model. Note that we have an additional complication here: only cells that are on the surface are able to replicate and migrate, and if these processes are stochastic, it is possible that a random fluctuation causes a population of mutant cells to disappear from the surface. For example, if the surface is rough and fluctuating, a mutant strain can be “swallowed” by fluctuations in the growing surface if it does not “surf” on the expanding frontier of the lesion and fails to establish a macroscopic sector fast enough [46, 47]. We thus need to consider the “surfing probability” *P*_surf_ that if a mutant was generated at time (lesion age) *a*_0_, it is still on the surface at time *a*.

Let us focus on a specific example of surface growth: Eden-like growth from Ref. [12]. It has been shown [48] that the probability *P*_surf_ that a mutant strain remains on the surface is approximately *P*_surf_ ≈ *c*/*a*_0_, i.e., inversely proportional to the age at which the first mutant cell has been created. This inverse proportionality law is likely to be true for more general models that fall into the KPZ universality class [49]. The proportionality constant c can be thought of as a “surfing time”, and while we are not aware of an *ab initio* estimate, our simulations indicate that it is on the order of 1/*v*_*n*_ days for the model considered in Ref. [12]. Since this constant depends on the microscopic details of the model, for the sake of generality we shall not ascribe a specific value in the formulas below.

New mutants emerge in this model with a rate equal to the mutation probability *p*_*µ*_ times the rate of expansion of the surface, which for sufficiently large (and old) lesions of age *a* will scale as *a*^2^. Following Ref. [22], we calculate the probability *r*_*n*_ that a randomly selected surface cell is non-mutant at radius *ρ* as

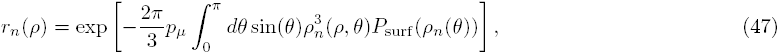

where *ρ*_*n*_(*ρ,θ*) = *ρe*^−|*θ*|/*βn*^ and 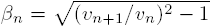 (c.f. equations (19),(23), and (25) in [22]). The formula (47) gives the fraction of type-*n* cells, but as a function of the radius rather than the age *a*. Equation (47) also accounts for mutant cells that remained on the surface, hence the factor *P*_surf_. By inserting *r* = *v*_*n*_*a* and *P*_surf_ = *c/a* we easily obtain that

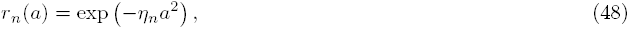

For

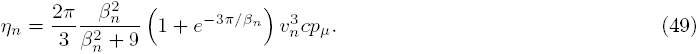

Note that if *v*_*n*+1_ = *v*_*n*_ and there is no selective advantage for additional mutations, then *η*_*n*_ vanishes, despite the fact that neutral mutants should have a non-zero chance of fixing: this model therefore does not apply to the stochastic fixation of neutral mutations.

For sufficiently low mutation rates and short times (namely, *η*_*n*_*a*^2^ ≪ 1) we can approximate Eq. (48) as

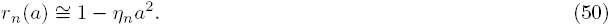

Equation (50) is quadratic in *a* and leads to an analytically solvable model, see the Supplementary Mathematica notebook. Notice that the fraction *r*_*n*_(*a*) can become negative for 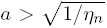 which limits the applicability of (50) to times shorter than 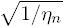. This problem can be partially alleviated by including higher order terms in the Taylor expansion of Eq. (48) at the expense of increasing calculation time.

### Comparison to simulations

We compared the results obtained in previous sections with numerical simulations of two models: our model as defined in section 1 in which growth and mutations are deterministic but migration of cells is a stochastic process, and the Eden-like lattice model [12] in which all these processes are stochastic.

### A. Computer simulations of the original model

We performed numerical simulations of the surface growth model. We treated individual lesions as balls of volume (4*π*/3)(*v*_*n*_*a*)^3^, where *a* was the age of the lesion. New lesions were established with rate 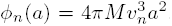.A new lesion was assigned type *n* with probability *r*_*n*_(*a*) = exp(−*µa*), and type *n* +1 with probability 1− *r*_*n*_(*a*). We assumed that mutations had the same effects on growth as in *Driver mutations with decreasing selective advantage:* three strong drivers and a small fitness advantage *ϵ* for each additional driver after the first three. The expansion speeds *v*_*n*_ were thus given by Eqs. (42-44).

In each simulation we measured the total volume, the average number of drivers per cell 〈*n*〉, and the proportion of cells with a specific number of mutations *n*, and then averaged those quantities over 500 such simulations. Fig. 4 shows an excellent agreement between simulations and the analytic formulae at early times; a small deviation can be seen for longer times. This deviation is caused by the analytic calculations neglecting stochastic effects and reproducing only “mean” behaviour.

The impact of stochastic fluctuations can be best seen in Fig. 4C which shows the average number of drivers 〈*n*〉 as a function of time *t*. If we first calculate the average number of drivers in each replicate simulation, and then average over simulations (“Method 1”), the obtained 〈*n*〉 deviates significantly from the analytical curve (purple points in Fig. 4C). However, if we calculate volumes of cells of type *V*_*n*_ for each simulation, average this over replicates, and then calculate 〈*n*〉 = Σ_*n*_ *n* 〈*V*_*n*_〉 / 〈*V*_tot_〉 (“Method 2”), we obtain a much better agreement (black points in Fig. 4C). The second method corresponds to what we do in the analytic calculations - we average out randomness already in Eqs. (1, 2) and use the average volumes to find 〈*n*〉. The discrepancy between Method 1 and the analytical calculations is caused by a broad and highly skewed distribution of the times by which lesions with new drivers arise in the stochastic simulation. Tumours of different sizes are weighted equally in Method 1, and hence the contribution to 〈*n*〉 from tumours in which drivers arose late is significant, even though such tumours are much smaller. These tumours will however decrease 〈*n*〉 which is exactly what we see in Fig. 4C.

### B. Comparison to the Eden lattice model

In the lattice model [12], each site is either unoccupied (which may be interpreted as being occupied by a normal cell) or occupied by a cancer cell. Each cell carries some number of driver mutations *n* ≥ 1. The simulation begins with a single cell with one driver mutation situated at the origin. Each time step, cells attempt to divide and produce one additional cell at a random lattice site which neighbours their own with rate *b*_*n*_, but the attempt is only successful if the random lattice site is empty. During replication cells can gain additional drivers with probability *p*_*µ*_ per daughter cell. Cells also migrate and create new microlesions with rate *M*. Distinct microlesions are approximately spherical, and do not interact with one another.

In that the resulting cloud of microlesions are asymptotically linearly expanding spheres and migration and mutation are both stochastic, the model is qualitatively similar to our mathematical model, but has much more complicated microscopic dynamics. Moreover, in contrast to the original model considered in this work, replication and mutation are also stochastic processes. It is thus not obvious *a priori* that the analytical model should be a good approximation to the lattice model.

We first tested whether the fraction of mutated cells 1 − *r*_*n*_(*a*) agrees with Eq. (50) from Section *More realistic surface fractions r_n_*(*a*). Figure 5 shows that the average 1 – *r*_1_(*a*) obtained from computer simulations is very close to *η*_1_*a*^2^, with *η*_1_ fitted to data from simulations of a single tumour. The distribution of 1 − *r*_1_(*a*) for a given *a* is however very broad. We thus conclude that our analytic model should be able to reproduce the average *V*_*n*_(*t*) from computer simulations, although stochastic effects may be visible in 〈*n*〉.

**Figure 5:**
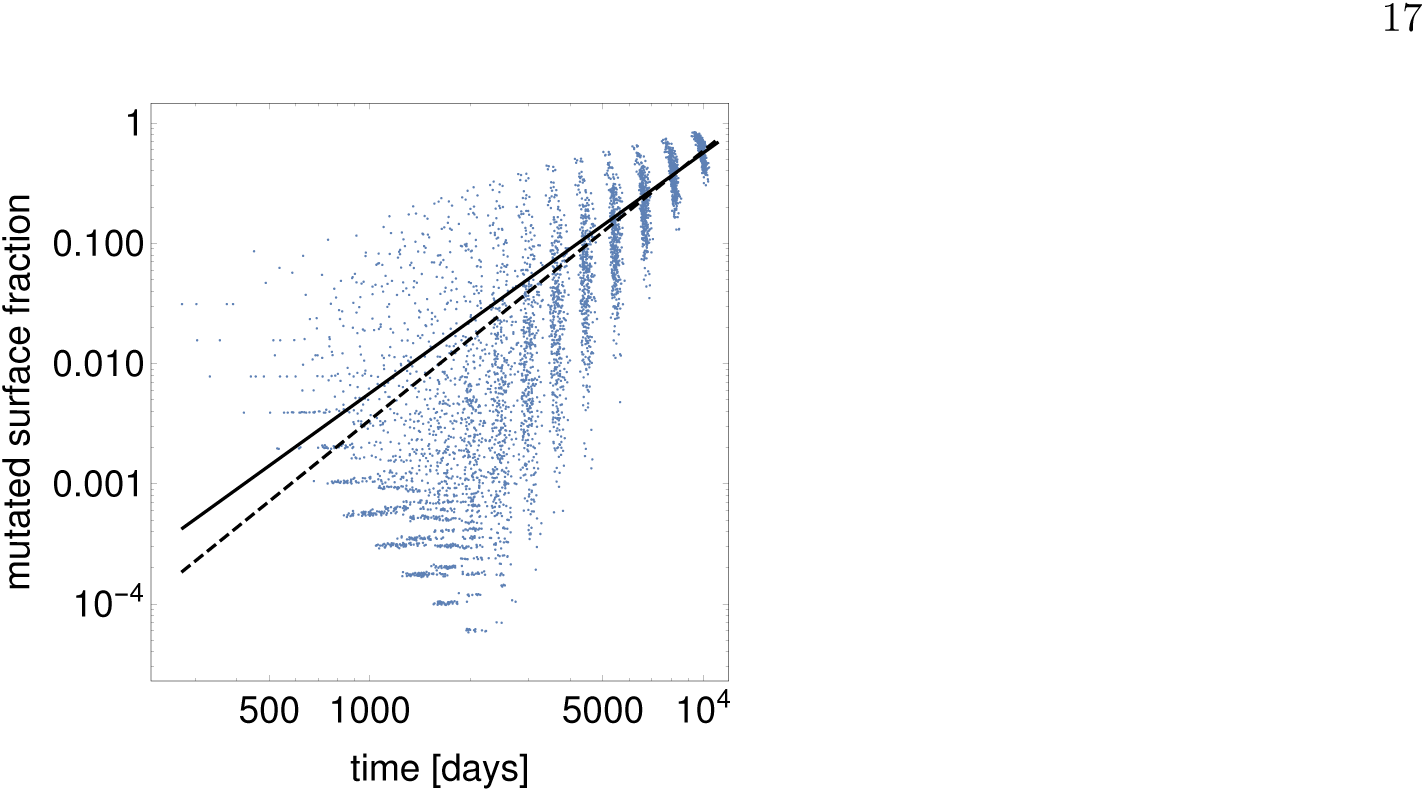
The fraction of mutants 1 − *r*_1_ on the surface of a single ball of cells in the Eden lattice model (blue points), compared to the quadratic approximation 1 − *r*_1_ = *η*_1_*a*^2^ (solid black line) and 1 − *r*_1_ = *Ca*^*γ*^ (dashed black line), with *η*_1_, *C*, *γ* fitted to the data points. The parameter values for this simulation are *b*_*1*_ = 0.0182 day^−1^, *M* = 0, *p*_*µ*_ = 10^−4^, and *ϵ* = 0.2.

We then compared the computer model with the analytical model. We assumed (similarly to Eqs. (42-44)) only three drivers:

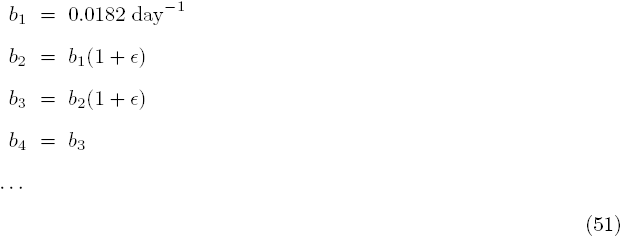

in the computer Eden-like model and, accordingly, equations (42-44) with *v*_1_ ≈ 0.53*b*_1_ [sites/day] in the mathematical model. The proportionality constant 0.53 was obtained by fitting *V*_tot_ (Eq. (19) in the analytical model) to the simulation data for *p*_*μ*_ = 0, using *G*_1_ as the single fitting parameter (see supplementary Mathematica notebook for further details).

We finally simulated the model with *p*_*µ*_ > 0, see Fig. 6. Since we did not know the exact relationship between the mutation rate and the selective advantage in the Eden model, and the corresponding parameters *p*_*µ*_, *ϵ* in the analytical model, we fixed *η*_*n*_ in Eq. (50) by fitting the product *cp*_*µ*_ from Eq. (49) to the numerical simulation data for the total volume *V*_tot_(*t*) (Fig. 6A). We then used the fitted parameters to calculate the average number of drivers. Figure 6B shows that the agreement between the two models is quite good even though the analytical model has vastly simpler dynamics than the Eden model. The analytical model slightly underestimates the number of driver mutations, and we attribute this to a broad distribution of times at which drivers first appear, as in Sec. *Computer simulations of the original model*.

**Figure 6:**
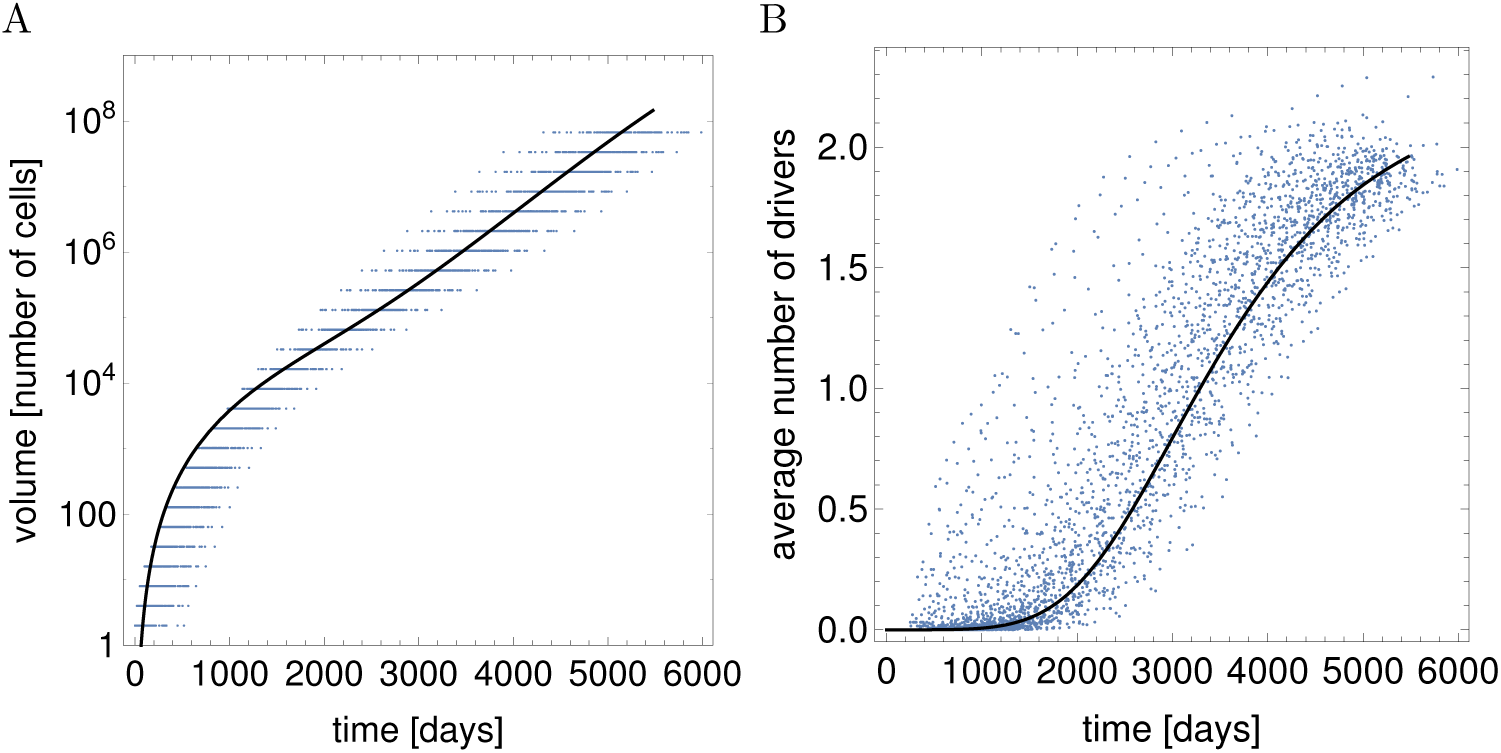
Comparison between the mathematical model and the Eden lattice model. Continuous lines are analytic solutions of the model, points correspond to computer simulations. A: Total volume as a function of time. B: The number of drivers versus time. Parameters for the simulations are *b*_1_ = 0.0182 day^−1^ and *n*_max_ = 3: for the analytics, *r*_*n*_ is given by Eq. (50) with *c* fitted to the data, *v*_*n*_ ≈ 0.53*b*_*n*_ = 0.00964 day^−1^. Both simulations and analytic calculations assume *M* = 10^−4^, *ϵ* = 0.5, and *p*_*µ*_ = 10^−4^.

### Discussion

In this paper we study a model of cancer based on the following three processes: replication of cancer cells, mutations endowing cells with fitness advantage, and migration that causes cells to disperse. The latter process causes the tumour to become a conglomerate of microlesions. Cellular dispersal has been recently recognized as a method by which tumours [28, 50] or, more generally, populations of motile cells [51] can speed up their growth. Our model applies both to local migration as well as long-range invasion involved in metastasis and is one of hallmarks of cancer [52]. In fact, the aforementioned conglomerate of lesions can also include metastatic lesions.

The strength of the model presented in this work lies in the relative ease with which its average behaviour can be obtained analytically. Perhaps surprisingly, including dispersal of cells in the model makes it easier to solve than most other spatial (non-well mixed) models, because migration effectively “smears out” spatial structure, bringing the model’s qualitative behaviour closer to well-mixed models. Analytical solubility means that the model works for tumours of any size, including large masses that need to be surgically removed, and it can be thus used to model cancer progression in humans. Below we discuss the most important implications of the model. These predictions deal with two aspects of cancer: growth laws, and genetic heterogeneity of tumours.

#### Tumour growth

In the absence of new driver mutations and assuming sufficient migration, our model predicts that long-time growth is exponential. This is also true when individual microlesions grow sub-exponentially, and even if their growth slows down over time. Given that most tumours contain avascular areas where the lack of oxygen and glucose inhibits proliferation [53, 54], our model provides a plausible explanation how the growth of an entire tumour can still be exponential, as often observed experimentally for intermediate-size tumours. This phenomenon does not require postulating any previously unknown mechanisms, but it relies on short-range migration of cancer cells, a process that certainly occur in nature and has been actively researched recently [55].

Figure 7A shows a region in the space of parameters *M*, *v*_1_ (*v*_1_ ≡ *v* in this case) for which the model predicts the total size to fall between 10^10^ and 10^12^ cells after 10 to 20 years - typical sizes of cancerous tumours after that time as explained earlier. Generally, the slower individual microlesions grow, the larger the migration probability needs to be to achieve the typical size, and the region of “good” parameters is approximately a straight line on the log-log plot of *M* versus *v*: *M* ~ *v*^−3^. This follows from how these parameters determine the growth rate *G* ~ (*Mv*^3^)^1/3^ and is related to the assumed surface growth of tumour microlesions in 3d space. If growth slows down over time (as studied in detail in “Surface growth with decreasing replication rate”), the “good” region moves up to higher *M* and *v*_1_ but it also becomes thinner. Assuming that growth occurs in a layer that is about 10 cell thick [53], and that cells replicate with rate *b* = 1 day^−1^, we can estimate that a selective advantage of the initial driver (b − *d*)/*b* = 0.001,…, 0.01 [8] correspond to *v*_1_ ≈ 10(b − d) = 0.01,…, 0.1. From figure 7A we can then deduce that the required migration rate *M* ≈ 10^−6^ – 10^−3^ - a rather modest number given that neoplastic cells can be highly motile [56, 57]. Note that the model would not be appropriate for very large selective advantages (~ 50% has been claimed for the first driver in colorectal cancer [58]) because *M* would have to be less than 10^−10^ in which case migration would be negligible even for macroscopic lesions. Our model is thus applicable to clonal expansions occurring after the initial driver has been acquired, because subsequent driver mutations are likely to be less potent.

**Figure 7:**
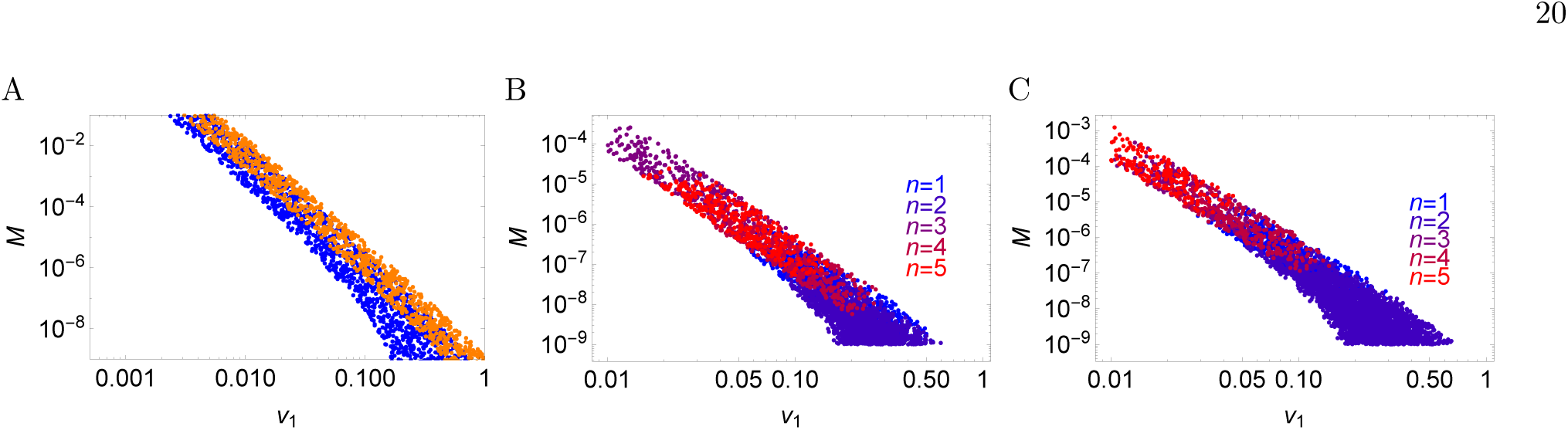
Regions in the space of parameters *M* and *v*_1_ which give *V*_tot_ between 10^10^ and 10^12^ when the tumour is 10…20 years old. To obtain these plots, 10^5^ points in the space (*v*_1_, *M*) were sampled and the volume of the tumour was calculated using the exact formulas. A: single-type model (here *v*_1_ ≡ *v*) with surface growth (blue) and slow-down growth (orange). B,C: multiple-types surface growth model with additional restriction that the average number of drivers is below 6 (including the initial driver *n* = 1). Colours correspond to different number of drivers. *µ* = 10^−5^ in panel B and *µ* = 10^−6^ in panel C. The region in which multiple drivers accumulate in the tumour shifts towards smaller *v*_1_/larger *M* as *µ* decreases.

If new driver mutations, each with a non-zero fitness advantage, steadily accumulate during tumour growth, our model predicts faster-than-exponential growth. While this speed-up does not have to be large and may arise only in tumours that are too big to be clinically significant, we are not aware of any evidence that super-exponential growth has been observed in human cancer. On the other hand, if only a few drivers can occur, growth will be exponential even at large times, which is consistent with the experimental evidence. We shall come back to this problem when discussing tumour heterogeneity.

Exponential or faster growth is obviously unrealistic for very large tumours for which spatial constraints become important. Many models of tumour growth have been proposed in the past [59] that account for the experimentally observed sigmoidal growth curve of many tumours. Many of these models are however phenomenological and are not based on the microscopic dynamics of tumour cells, in contrast to the model studied here. It would be interesting to learn what minimal changes our model would require to reproduce sigmoidal growth.

#### Genetic heterogeneity of tumours

Experimental evidence gathered over the last 6 years [10, 60, 61] strongly suggest that cancerous masses are genetically heterogeneous, although the exact level of heterogeneity and its importance are still under debate. In this work we explored this heterogeneity by calculating the size of clonal subpopulations of cells with different numbers of driver mutations. We showed that these subpopulations increase exponentially in size and that only one driver is initially dominant, until finally becoming replaced by a mixture of clones with two, three, and more drivers. The coexistence of multiple clones means that the time to *n* drivers cannot be simply calculated as the sum of the times between consecutive driver mutations.

We have also shown that if each new driver increases the selective advantage of cancer cells in comparison to normal cells, the average number of driver mutations predicted by our model increases exponentially in time for all considered scenarios. This means that most drivers would accumulate late during cancer progression. There is limited evidence [45, 61] that this is not true. On the other hand, if only a few first drivers have significant fitness advantage, these drivers will accumulate early during growth and the tumour will become much more homogeneous. An interesting application of our model would be to predict how strong the selective advantage of new drivers can be that is still consistent with recently postulated neutral evolution in tumours [61].

We can also use the model to infer the relationship between the number of drivers and the parameters of the model for “typical” tumours. Figure 7B, shows a “good” region in (*M*, *v*_1_) which produce “typical” tumour sizes. Points in the region have been colour-coded depending on how many drivers the tumour has for a given pair (*M*, *v*_1_). The plot shows that, unless individual microlesions grow slowly enough (small *v*_1_), the number of new drivers is close to zero (only one, initial driver present). Since *v*_1_ is related to the selective advantage of a driver, this is consistent with previous research [8] showing that significant accumulation of new drivers occurs only for small selective advantages.

Our model can be extended in several ways. For example, we have not analysed spatial distribution of drivers. While it may not be possible to do this analytically, the model proposed here can easily be simulated on a computer (as demonstrated in Section “Comparison to simulations”) and perhaps extended to include spatial locations of microlesions. Another possible extension would be to consider that only a fraction of cells in the tumour (“cancer stem cells”) can replicate forever, whereas the majority of cells can undergo only a few rounds of replication, as e.g. in Ref. [62]. This assumption would affect the growth rate and the rate at which mutations accumulate in a single lesion. Finally, since separability of the coupled systems of structured population equations (1,2) only depends on the validity of the infinite genome approximation, it is likely that our approach is extensible to evolution in more complex landscapes than the linear chain of mutations we have studied. In particular, the model could be extended to epistatic interactions between drivers and passangers [63] or non-equal, random fitness increments [64].

## Acknowledgments

We thank EPSRC for support. B.W. was supported by a Scottish Government/Royal Society of Edinburgh (RSE) Personal Research Fellowship.

## Author Contributions

C.P., M.A.N., and B.W. conceived the model. C.P. derived the analytical results. C.P. and B.W. made the figures. All authors wrote the manuscript.

## Additional Information

*Competing financial interests:* The authors declare no competing financial interests.

## Methods

### General solution with multiple types of microlesions

We first consider the general case (1,2). Given the initial condition *f*_*n*_(*a*, 0) = *δ*_*n*,1_*δ* (*a*) i.e. that there is only one lesion of type *n* = 1 and age zero at time *t* = 0, Eq. (1) implies that *f*_*n*_(*a*, *t*) takes the form

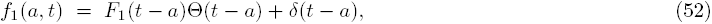

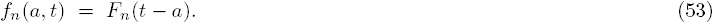

The boundary condition (2) can be rewritten as

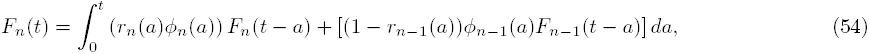

which upon a Laplace transform and little algebra gives

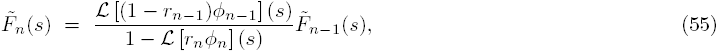

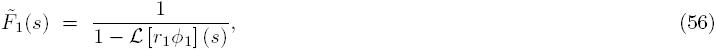

where ℒ[*…*](*s*) denotes the Laplace transform of the function in the square brackets. We can now write a formal solution of our coupled system of equations for *n* > 1:

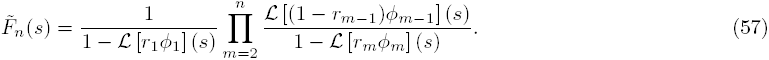

This clearly has simple poles at every *G*_*m*_ where

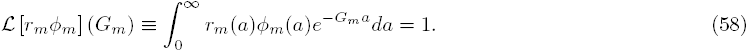

which is the Euler-Lotka equation for each lesion type *m*. In the large-*t* limit, the number of microlesions of type *n* will thus grow exponentially, and the rate of this exponential growth will be the largest of all dominant roots *G*_*j*_ for 1 ≤ *j* ≤ *n*. In particular, if each new driver increases the product *r*_*m*_(*a*)*ϕ*_*m*_(*a*) (as if increasing the replication rate of cells in the lesion), the number of microlesions of type *n* will increase exponentially with rate *G*_*n*_, where *G*_*n*_ is the largest root of Eq. (58).

### Surface growth with many drivers

The required Laplace transforms read

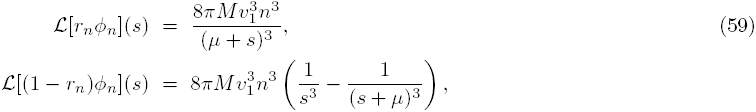

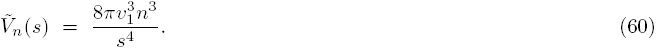

From Eqs. (58) and (59) we then find the exponential growth rate of the number of microlesions of type *n* (Eq. (33).

#### Number of lesions of type *n*

Upon inserting Eqs. (59,60) into Eq. (57) we obtain an expression for the Laplace transform 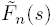. This expression can be analytically inverted to find *F*_*n*_(*z*) for *n* = 1, 2, 3,…, although the formulas for even the lowest *n* are quite complicated (SI Mathematica notebook). Nevertheless, all *F*_*n*_(*z*) take the form

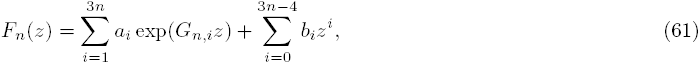

where *G*_*n*,*i*_ is the *i*-th root of Eq. (58) for a given *n*, and *a*_*i*_,*b*_*i*_ depend on *v*_1_,*M,µ* The number of microlesions of type *n* is thus

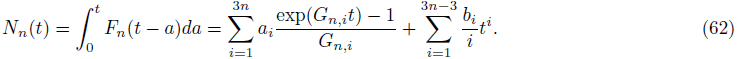

Similarly, the total volume (number of cells) of type *n* is

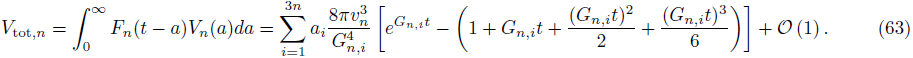

#### Asymptotic number of drivers

Let us consider the behaviour of 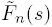 around *s* = *G_n_.* From Eq. (57) we obtain that 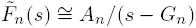 where

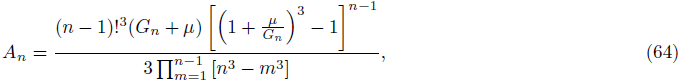

from which we obtain that *F*_*n*_(*z*) ≅ *A*_*n*_ exp(*G*_*n*_*z*). Let us now introduce the generating function

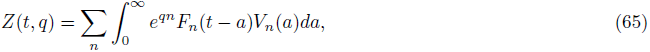

from which the average number of drivers can be calculated as:

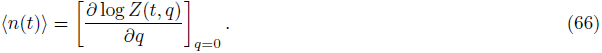

In the biologically-relevant limit, the mutation rate *µ* is considerably less than the net growth rate *G*_1_. Taking the limit *µ*/*G*_1_ → 0 we obtain

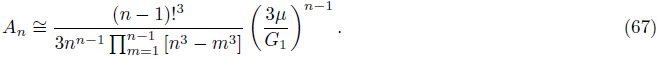

This enables us to write

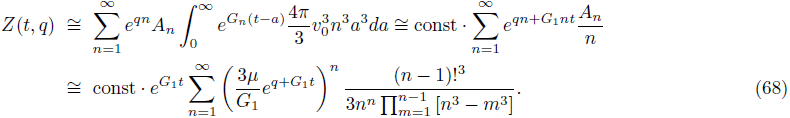

In the limit of large *n* we can approximate the complicated numerical factor in the above equation by

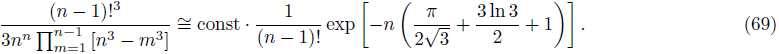

From Eq. (68) we then obtain that

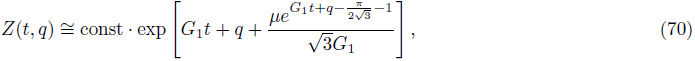

and finally, using Eq. (66), we obtain Eq. (35). We can also use Eq. (70) to calculate the asymptotic volume of the tumour,

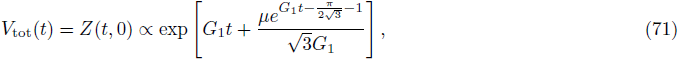

which shows the volume increases faster than exponentially over time due to the accumulation of driver mutations.

#### Slow-down surface growth

The Laplace-transformed distribution 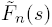 now reads

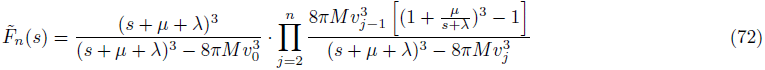

so that the exponential growth rate of the population of microlesions of type *n* is given by Eq. (36).

The coefficients *A*_*n*_, equivalent of Eq. (64), are

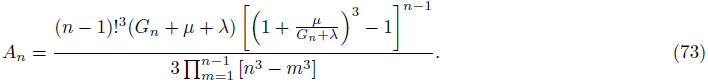

Following the same procedure as in the earlier section “Surface growth”, we can approximate *A*_*n*_ in the limit *µ*/*G*_1_ → 0 as

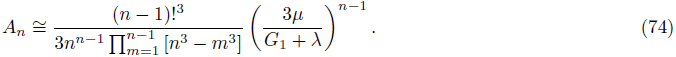

This gives

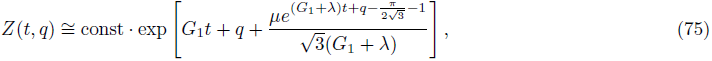

from which we can deduce, using Eq. (66), that the average number of drivers is given by Eq. (37).

### Volumetric growth

Equation (57) implies that

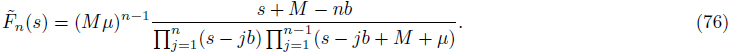

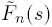 has poles at every *s* = *kb* for 1 ≤ *k ≤ n* and *s* = *kb* − *M* − *µ* for every 1 ≤ *k* ≤ *n* − 1. Consequently, *F*_*n*_(*z*) can be expressed exactly as

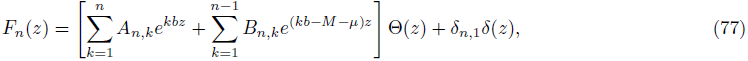

which leads to the following expression for the total volume of cells of type *n*:

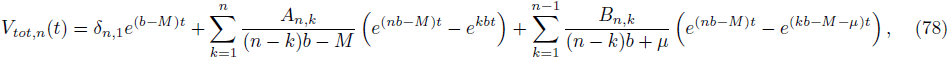

with the coefficients

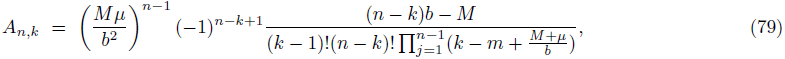

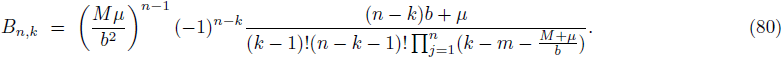

The above formulas are exact but quite complex. To obtain simpler expressions for the total volume and the number of drivers we notice that in real tumours *b* ≫ *M*, i.e., the replication rate is much larger than the migration rate, otherwise cells would only move around without replicating. The rate *µ* at which drivers occur is also much smaller than *b* [8]. Consequently, *F*_*n*_(*z*) is dominated by the term proportional to *A*_*n*,*n*_ at sufficiently long times (*bt* ≫ 1). We can also approximate *A*_*n*,*n*_ as

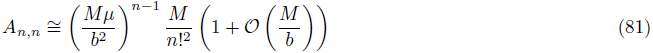

and the function *Z*(*t*, *q*) may be written as

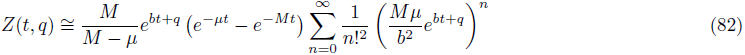

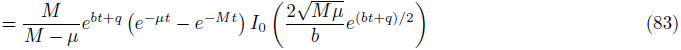

where *I*_0_(*x*) is a modified Bessel function of the first kind. This allows us to write immediately that

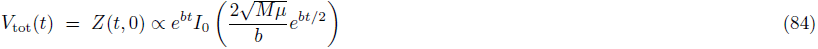

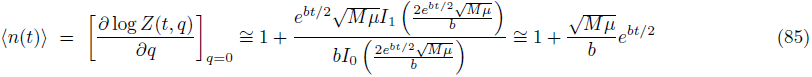

which is Eqs. (41).

### Numerical integration of equations (1,2)

As a check-up of our analytic calculations, the system of equations (1) with boundary conditions (2) were numerically integrated as follows. Time was discretized in steps of length Δ*t* days. We usually set Δ*t* to 1 day, and checked that making Δ*t* smaller did not visibly affect our results. The age distribution *f*_*n*_(*a,t*) was discretized as *f*_*n,i,t*_, where *i* = *a*/Δ*t*. The array of *f*_*n,i,t*_ was truncated at *i* = *t*_max_/Δ*t*, where *t*_max_ was set to 15 years. We also fixed the maximal possible number of drivers to *n*_max_ = 32.

We then discretized Eqs. (1,2) as

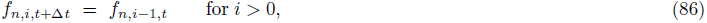

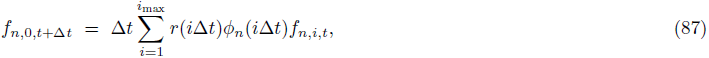

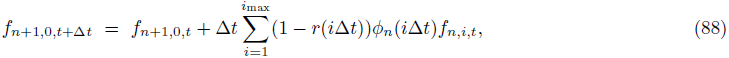

where the first equation corresponds to Eq. (1) shifting the distribution forward in time, and the remaining two equations account for the boundary condition (2). The volume of all microlesions of a given type does not enter into the dynamics explicitly, as is apparent from the form of (2), but it appears in the formulas for total tumour volume and mean number of drivers 〈*n*(*t*)〉:

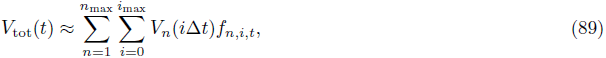

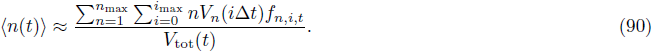

The computer program outputted *V*_tot_(*t*) and 〈*n*(*t*)〉 every 10Δ*t* = 10 days.

## References

[1] Tomasetti, C. & Vogelstein, B. Variation in cancer risk among tissues can be explained by the number of stem cell divisions. Science 347, 78–81 (2015).

[2] Sjöblom, T. et al. The consensus coding sequences of human breast and colorectal cancers. Science 314, 268–274 (2006).

[3] Wood, L. D. et al. The genomic landscapes of human breast and colorectal cancers. Science 318, 1108–1113 (2007).

[4] Jones, S. et al. Comparative lesion sequencing provides insights into tumor evolution. Proceedings of the National Academy of Sciences 105, 4283–4288 (2008).

[5] Vogelstein, B. et al. Cancer genome landscapes. Science 339, 1546–58 (2013).

[6] Tomasetti, C., Marchionni, L., Nowak, M. A., Parmigiani, G. & Vogelstein, B. Only three driver gene mutations are required for the development of lung and colorectal cancers. Proceedings of the National Academy of Sciences 112, 118–123 (2015).

[7] Armitage, P. & Doll, R. The age distribution of cancer and a multi-stage theory of carcinogenesis. British Journal of Cancer 8, 1 (1954).

[8] Bozic, I. et al. Accumulation of driver and passenger mutations during tumor progression. Proceedings of the National Academy of Sciences 107, 18545–18550 (2010).

[9] Makohon-Moore, A. P. et al. Clonal evolution defines the natural history of metastatic pancreatic cancer. Cancer Research 75, 4137–4137 (2015).

[10] Yachida, S. et al. Distant metastasis occurs late during the genetic evolution of pancreatic cancer. Nature 467, 1114–1117 (2010).

[11] Diaz Jr, L. A. et al. The molecular evolution of acquired resistance to targeted EGFR blockade in colorectal cancers. Nature 486, 537–540 (2012).

[12] Waclaw, B. et al. A spatial model predicts that dispersal and cell turnover limit intratumour heterogeneity. Nature 525, 261–264 (2015).

[13] Nowak, M. A. et al. The role of chromosomal instability in tumor initiation. Proceedings of the National Academy of Sciences 99, 16226–16231 (2002).

[14] Michor, F., Iwasa, Y., Lengauer, C. & Nowak, M. A. Dynamics of colorectal cancer. Seminars in Cancer Biology 15, 484–493 (2005).

[15] Dingli, D., Traulsen, A., Lenaerts, T. & Pacheco, J. M. Evolutionary Dynamics of Chronic Myeloid Leukemia. Genes & Cancer 1, 309–315 (2010).

[16] Wodarz, D. & Komarova, N. L. Dynamics of Cancer: Mathematical Foundations of Oncology (World Scientific, 2014).

[17] Altrock, P. M., Liu, L. L. & Michor, F. The mathematics of cancer: integrating quantitative models. Nature Reviews Cancer 15, 730–745 (2015).

[18] Iwasa, Y. & Michor, F. Evolutionary Dynamics of Intratumor Heterogeneity. PLoS ONE 6, e17866 (2011).

[19] Komarova, N. Stochastic modeling of drug resistance in cancer. Journal of Theoretical Biology 239, 351–366 (2006).

[20] Durrett, R., Foo, J., Leder, K., Mayberry, J. & Michor, F. Intratumor Heterogeneity in Evolutionary Models of Tumor Progression. Genetics 188, 461–477 (2011).

[21] Durrett, R. Population genetics of neutral mutations in exponentially growing cancer cell populations. The Annals of Applied Probability 23, 230–250 (2013).

[22] Antal, T., Krapivsky, P. L. & Nowak, M. A. Spatial evolution of tumors with successive driver mutations. Physical Review E 92 (2015).

[23] Nowak, M. A., Michor, F. & Iwasa, Y. The linear process of somatic evolution. Proceedings of the National Academy of Sciences 100, 14966–14969 (2003).

[24] Komarova, N. L. Spatial Stochastic Models for Cancer Initiation and Progression. Bulletin of Mathematical Biology 68, 1573–1599 (2006).

[25] Durrett, R., Foo, J. & Leder, K. Spatial Moran models II. Tumor growth and progression (2012).

[26] Durrett, R. & Moseley, S. Spatial Moran models I? Stochastic tunneling in the neutral case. The annals of applied probability: an official journal of the Institute of Mathematical Statistics 25, 104 (2015).

[27] Foo, J., Leder, K. & Ryser, M. D. Multifocality and recurrence risk: A quantitative model of field cancerization. Journal of Theoretical Biology 355, 170–184 (2014).

[28] Enderling, H., Hlatky, L. & Hahnfeldt, P. Migration rules: tumours are conglomerates of self-metastases. British Journal of Cancer 100, 1917–1925 (2009).

[29] Hanin, L., & Zaider, M. A stochastic model for the sizes of detectable metastases. Journal of Theoretical Biology 243(3), 407–417 (2006).

[30] StefanieJefrey, M., Carlson, R. W. & Stockdale, F. E. The importance of the lumpectomy surgical margin status in long term results of breast conservation. Cancer 76, 259–67 (1995).

[31] Suzuoki, M. et al. Impact of caveolin-1 expression on prognosis of pancreatic ductal adenocarcinoma. British Journal of Cancer 87, 1140–1144 (2002).

[32] McDonald, O. G., Wu, H., Timp, W., Doi, A. & Feinberg, A. P. Genome-scale epigenetic reprogramming during epithelial-to-mesenchymal transition. Nature Structural & Molecular Biology 18, 867–874 (2011).

[33] Drasdo, D. Buckling instabilities of one-layered growing tissues. Physical Review Letters 84, 4244 (2000).

[34] Montel, F. et al. Stress clamp experiments on multicellular tumor spheroids. Physical Review Letters 107, 188102 (2011).

[35] Iwata, K., Kawasaki, K. & Shigesada, N. A dynamical model for the growth and size distribution of multiple metastatic tumors. Journal of Theoretical Biology 203, 177–186 (2000).

[36] Michor, F., Nowak, M. A. & Iwasa, Y. Stochastic dynamics of metastasis formation. Journal of Theoretical Biology 240, 521–530 (2006).

[37] Keyfitz, B. L. & Keyfitz, N. The mckendrick partial differential equation and its uses in epidemiology and population study. Mathematical and Computer Modelling 26, 1–9 (1997).

[38] Perthame, B. Transport equations in biology (Springer, 2006).

[39] Baratchart, E. et al. Computational modelling of metastasis development in renal cell carcinoma. PLoS Comput Biol 11, e1004626 (2015).

[40] Folkman, J. What is the evidence that tumors are angiogenesis dependent? Journal of the National Cancer Institute 82, 4–7 (1990).

[41] Naumov, G. N., Folkman, J. & Straume, O. Tumor dormancy due to failure of angiogenesis: role of the microenvironment. Clinical & Experimental Metastasis 26, 51–60 (2008).

[42] Weinberg, R. A. The Biology of Cancer (Garland Science, 2007).

[43] Lavrentovich, M. O. & Nelson, D. R. Survival probabilities at spherical frontiers. Theoretical Population Biology 102, 26–39 (2015).

[44] Tomasetti, C., Marchionni, L., Nowak, M. A., Parmigiani, G. Vogelstein, B. Only three driver gene mutations are required for the development of lung and colorectal cancers. Proceedings of the National Academy of Sciences 112, 118–123 (2015).

[45] Sottoriva, A. et al. A Big Bang model of human colorectal tumor growth. Nat. Genet. 47, 209–216 (2015).

[46] Hallatschek, O., Hersen, P., Ramanathan, S. Nelson, D.R.. Genetic drift at expanding frontiers promotes gene segregation. Proceedings of the National Academy of Sciences, 104, 50, 19926–19930, (2007).

[47] Gralka, M. et al. Allele surfing promotes microbial adaptation from standing variation. Ecology Letters e12625 (2016). doi:10.1111/ele.12625

[48] Lavrentovich, M.O. Nelson, D.R.. Survival probabilities at spherical frontiers. Theoretical Population Biology 102, 26–39, (2015).

[49] Alves, S. G. Ferreira, S. C. Eden clusters in three dimensions and the Kardar-Parisi-Zhang universality class. Journal of Statistical Mechanics: Theory and Experiment 2012, P10011 (2012).

[50] Thalhauser, C. J., Lowengrub, J. S., Stupack, D. & Komarova, N. L. Selection in spatial stochastic models of cancer: Migration as a key modulator of fitness. Biol. Direct 5, 21 (2010).

[51] Hallatschek, O. & Fisher, D. S. Acceleration of evolutionary spread by long-range dispersal. Proceedings of the National Academy of Sciences 111, E4911–E4919 (2014).

[52] Hanahan, D. & Weinberg, R. A. The Hallmarks of Cancer. Cell 100, 57–70 (2000).

[53] Vaupel, P., Kallinowski, F. & Okunieff, P. Blood Flow, Oxygen and Nutrient Supply, and Metabolic Microenvironment of Human Tumors: A Review. Cancer Research 49, 6449–6465 (1989).

[54] Parks, S. K., Cormerais, Y., Marchiq, I. & Pouyssegur, J. Hypoxia optimises tumour growth by controlling nutrient import and acidic metabolite export. Molecular Aspects of Medicine 47-48, 3–14 (2016).

[55] Fokkelman, M. et al., Cellular adhesome screen identifies critical modulators of focal adhesion dynamics, cellular traction forces and cell migration behaviour. Scientific Reports 6, 31707 (2016).

[56] Enterline, H. T. & Coman, D. R. The ameboid motility of human and animal neoplastic cells. Cancer 3, 1033–1038 (1950).

[57] Friedl, P. & Wolf, K. Tumour-cell invasion and migration: diversity and escape mechanisms. Nature Reviews Cancer 3, 362–374 (2003).

[58] Vermeulen, L. et al. Defining Stem Cell Dynamics in Models of Intestinal Tumor Initiation. Science 342, 995–998 (2013).

[59] Rodriguez-Brenes, I. A., Komarova, N. L. & Wodarz, D. Tumor growth dynamics: insights into evolutionary processes. Trends in Ecology & Evolution 28, 597–604 (2013).

[60] Gerlinger, M. et al. Intratumor Heterogeneity and Branched Evolution Revealed by Multiregion Sequencing. New England Journal of Medicine 366, 883–892 (2012).

[61] Ling, S. et al. Extremely high genetic diversity in a single tumor points to prevalence of non-darwinian cell evolution. Proceedings of the National Academy of Sciences 112, E6496–E6505 (2015).

[62] Werner, B. et al. The cancer stem cell fraction in hierarchically organized tumors can be estimated using mathematical modeling and patient-specific treatment trajectories. Cancer Research 76, 1705 (2016).

[63] Bauer, B., Siebert, R. & Traulsen, A. Cancer initiation with epistatic interactions between driver and passenger mutations. Journal of Theoretical Biology 358, 52–60 (2014).

[64] Durrett, R., Foo, J., Leder, K., Mayberry, J. & Michor, F. Evolutionary dynamics of tumor progression with random fitness values. Theoretical Population Biology 78, 54–66 (2010).

